# Habitat resource overlap in two broad-ranged sympatric Neotropical forest eagles

**DOI:** 10.1101/2022.03.24.485595

**Authors:** Luke J. Sutton, David L. Anderson, Miguel Franco, Felipe Bittioli R. Gomes, Christopher J.W. McClure, Everton B.P. Miranda, F. Hernán Vargas, José de J. Vargas González, Robert Puschendorf

**Affiliations:** School of Biological and Marine Sciences, University of Plymouth, Drake Circus, Plymouth, PL4 8AA, United Kingdom; The Peregrine Fund, 5668 West Flying Hawk Lane, Boise, Idaho 83709 USA; Faculdade de Etnodiversidade, Educação do Campo; Programa de Pós-Graduação em Biodiversidade e Conservação – PPGBC; Universidade Federal do Pará, Campus Universitário de Altamira, Pará

**Keywords:** Crested Eagle, environmental ordination, habitat use, Harpy Eagle, resource partitioning, resource selection functions

## Abstract

Quantifying resource partitioning between co-occurring species has important ecological and evolutionary implications. Yet, few studies compare resource overlap in both geographic and environmental space. We test whether the habitat requirements of two closely related Neotropical forest eagles, the crested eagle (*Morphnus guianensis*) and harpy eagle (*Harpia harpyja*), differ at fine and coarse resolutions across their shared geographic range. Using landcover and topographic covariates, we quantified resource overlap first using higher resolution (30 arc-sec data) generalized linear models (GLMs), and second using coarser-grain (2.5 arc-min data) environmental ordination. The distribution of both eagles was largely explained by canopy species richness and structural complexity with evergreen forest, but with differing responses to landcover and topography, particularly with the harpy eagle more likely in areas of dense evergreen forest. Both eagles were negatively associated with mosaic forest, with this relationship stronger for the crested eagle. Harpy eagle distribution was restricted by higher elevation and terrain roughness, compared to the crested eagle, whose distribution was more restricted by canopy species richness and structure. From the GLMs, resource overlap was > 92 % in geographical space but reduced to 64 % in environmental space. From ordination, resource overlap was 76 % in environmental space, with randomization tests supporting equivalent environmental space for both eagles. Our results suggest that at the biogeographical scale, crested and harpy eagles share environmental space, but there may be subtle differences in fine-scale habitat preference. We recommend habitat resource overlap be assessed in both geographical and environmental space at multiple resolutions to capture the inherent variability in environmental conditions available to co-occurring species.

## Introduction

The distribution of specific habitat resources is a major factor determining species range limits at both fine and broad spatial scales (Morrison *et al*. 2006). However, species interactions are also important for influencing animal distributions (MacArthur 1972; Cody 1974). Whilst support for habitat as a key factor limiting animal distributions is evident (Dorazio *et al*. 2015; Veech 2021), habitat resources alone cannot fully explain a given species distribution (Thompson 2005; Anderson 2017), or where and how co-occurring species coexist when they have overlapping ranges (Diamond 1970; MacArthur 1972). In highly biodiverse tropical forests, closely related species may exhibit patchy, but overlapping distributions (MacArthur 1972; Robinson 1994). This may result in certain species only being able to co-exist through resource partitioning (Levins 1968; Cody 1974), where specific and subtle differences in resource and habitat requirements are met.

In Neotropical forests, raptors are amongst the least-studied bird groups due to their inherent rarity and large area requirements, which present difficulties working in remote, logistically challenging environments. Compared to temperate regions, current knowledge on the environmental drivers of distribution for many Neotropical raptors is limited (Buechley *et al*. 2019), despite the Neotropics being a raptor diversity hotspot (McClure *et al*. 2018). Among Neotropical forest raptors, the crested eagle (*Morphnus guianensis*) and harpy eagle (*Harpia harpyja*) have almost identical geographical ranges across Central and South America, occurring widely across lowland broadleaf rainforests (Ferguson-Lees & Christie 2005; Gomes & Sanaiotti 2015; Miranda *et al*. 2019; Sutton *et al*. 2021a,b). Both species are uncommon with low population densities, with the harpy eagle being on average ~1.3 times larger than the crested eagle (Ferguson-Lees & Christie 2005). Both eagles are monotypic and the only members of the subfamily Harpiinae in the Neotropics (Lerner & Mindell 2005; Brown & Amadon 1968).

Generally, the crested eagle feeds on smaller and more diverse prey than the harpy eagle, mainly arboreal snakes, small primates, opossums, and birds (Bierregaard 1984; Julliot 1994; Robinson 1994; Whitacre *et al*. 2012; Gomes *et al*. 2021), whereas harpy eagle diet is largely comprised of large arboreal mammals, primarily sloths and primates (Aguiar-Silva *et al*. 2014; Miranda 2015). Thus, crested and harpy eagles may be able to co-exist by partitioning food resources, avoiding direct competition (Sanaiotti *et al*. 2015). On a microhabitat scale, the two species may use differing forest canopy strata for breeding and hunting (Gomes *et al*. 2021), with harpy eagles nesting in canopy-emergent trees (Sanaiotti *et al*. 2015; Miranda *et al*. 2020) and crested eagles canopy trees (Bierregaard 1984; Whitacre *et al*. 2012). Observations of interspecific feeding of a fledged harpy eagle at its nest by a female crested eagle (Vargas González *et al*. 2006) and interactions at nest sites between the two species (Aguiar-Silva *et al*. 2019) suggest that territorial behaviour can be relaxed when both eagles exist in close proximity to each other.

Apart from localized observations reporting both Harpiinae eagles breeding within ~1 to 3 km of each other in similar habitats (Galetti *et al*. 1997; Muñiz-López *et al*. 2007; Sanaiotti *et al*. 2015), little is known about how these two eagles co-exist at a large biogeographical scale with highly overlapping geographic ranges. Both eagles were recorded in similar forest habitats from two landscape habitat selection studies, one in an 800-ha section of Amazonian Peru (Manu National Park; Robinson 1994), and the second along a 276 km stretch of the Xingu river in Amazonian Brazil (Sanaiotti *et al*. 2015). Harpy eagles were observed more frequently than crested eagles, suggesting the harpy eagle may be the more common of the two species where both are present. From surveys along the Xingu river, Sanaiotti *et al*. concluded that territories of both eagles overlapped, but this is contrary to earlier results from a 10,000-ha study area in French Guiana which suggested that territories of neither eagle overlapped (Thiollay 1989).

At a local level, both eagles may partition food resources and prefer different micro-habitats. However, an unanswered question is whether these patterns scale up to a broader biogeographical extent where both eagles may prefer differing habitat. Because both Harpiinae eagles co-occur within similar geographical ranges, the expectation in geographical space is for a high level of habitat resource overlap. However, given that habitat resources are rarely equally distributed across landscapes, geographical comparisons between sister species may over-represent the overlap in environmental requirements. Therefore, when assessed in environmental space, habitat resource overlap may be lower indicating environmental niche partitioning (Pulliam 2000; Bleyhl *et al*. 2018). Even so, given that narrow niche breadth is viewed as optimal in the relatively stable and predictable environment of the tropics (Levins 1968), both eagles are expected to have specialized habitat requirements irrespective of the level in habitat resource overlap.

Previous studies have reported differences in habitat use at localized scales for the crested and harpy eagle (Robinson 1994; Thiollay 1989; Sanaiotti *et al*. 2015), but how habitat differences limiting distribution are perceived at broader biogeographical scales is unknown. Here, we use a multi-scale assessment across the entire environmental and geographic space available to both eagles to quantify habitat resource overlap and preference for specific habitats. We use Resource Selection Functions (RSFs) to first determine the fine-grain habitat correlates of distribution in both geographical and environmental space across the entire shared range for both eagles. Second, we perform an environmental Principal Component Analysis (PCA-env) to quantify the level of coarse-grain resource overlap solely in environmental space as a comparison to using the finer-scale RSFs. Given their almost identical distributions across lowland tropical forest, we predict that, (1) the range limits of both eagles will be restricted by similar habitat types, and (2) both eagles will have specialized habitat requirements when quantified in both geographic and environmental space.

## Methods

### Eagle occurrence data

We sourced crested and harpy eagle occurrences from the Global Raptor Impact Network (GRIN, McClure *et al*. 2021) consisting of occurrence data from the Global Biodiversity Information Facility (GBIF 2019a,b), which are mostly eBird records (crested eagle = 75 %, harpy eagle = 79 %, Sullivan *et al*. 2009). Three additional occurrence datasets for both the crested eagle (Gomes & Sanaiotti 2015) and harpy eagle range (Vargas González & Vargas 2011; Miranda *et al*. 2019) were also incorporated into our dataset. We cleaned occurrences by removing duplicate records, and those with no geo-referenced location. Only occurrences recorded from year 2000 onwards were included to temporally match the habitat covariates. We quantified spatial overlap in the point occurrences using a point-proximity overlap metric (*O*; Cardillo & Warren 2016) to test if both sets of occurrence points for each eagle species were distributed randomly and independent from each other. The point-proximity *O* metric tests for spatial overlap based on the co-aggregation in the point occurrences for each respective eagle species. The metric ranges from zero to one, with a value ~0.5 expected if the occurrence points of both species have a random and independent distribution.

We compiled totals of 881 crested eagle records and 1065 harpy eagle records after data cleaning, with unfiltered occurrences having a moderately random distribution (*O* = 0.230, Fig. 1). We applied a 1-km spatial filter between each occurrence point to reduce sampling bias for the RSFs using the ‘geoThin’ function in the R package enmSdm (Smith 2019). Using a 1-km filter approximately matches the resolution of the raster data (~1-km) and reduces the effect of biased sampling (Kramer-Schadt *et al*. 2013). Applying the 1-km spatial filter resulted in 439 occurrences for the crested eagle and 801 occurrences for the harpy eagle. Spatially filtered occurrences for both eagle species were distributed randomly and independent from each other (*O* = 0.448) compared to the unfiltered occurrences.

**Figure 1.**
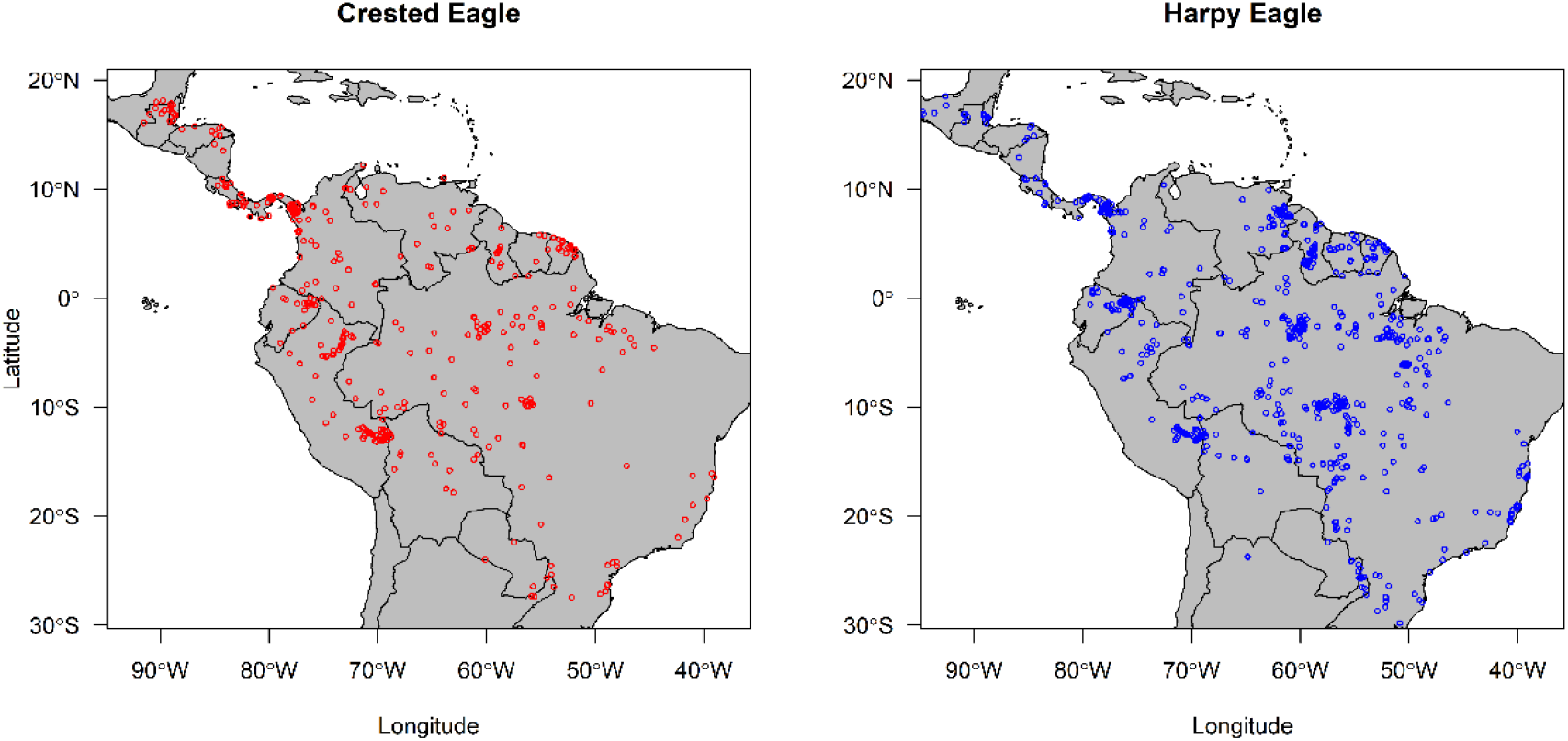
Distribution of crested eagle and harpy eagle occurrences across Central and South America sourced from the Global Raptor Impact Network (GRIN) data information system.

### Habitat covariates

Current distribution maps show both crested and harpy eagles exist in similar geographical space (Fig. 1, Ferguson-Less & Christie 2005), thus our expectation was for their respective distributions to be limited by the same habitat factors. Potential covariates representing topography, canopy vegetation heterogeneity, and landcover were downloaded from the ENVIREM (Title & Bemmels 2018) and EarthEnv (www.earthenv.org) databases. Biologically relevant covariates were selected *a prioiri* based on the key limiting habitat factors related theoretically and empirically to each species distribution in lowland tropical forests (Vargas González *et al*. 2006; Whitacre *et al*. 2012; Sanaiotti *et al*. 2015; Miranda *et al*. 2019). We included a total of six continuous covariates (Table 1) in the RSF analyses at a spatial resolution of 30 arc-sec (~1-km resolution). Raster layers were cropped using a delimited polygon consisting of all known range countries (including the states of Formosa, Jujuy, Misiones and Salta in northern Argentina, and the states of Chiapas, Oaxaca and Tabasco in southern Mexico), further improving model predictive power by reducing the background area used for testing points used in model evaluation (Radosavljevic & Anderson 2014).

**Table 1.** Habitat covariates used in all spatial and modelling analyses for the crested and harpy eagle.

| Covariate | Source | Citation |
| --- | --- | --- |
| Cultivated (%) | EarthEnv | Tuanmu & Jetz 2014 |
| Elevation (m) | EarthEnv | Amatulli *et al.* 2018 |
| Evergreen forest (%) | EarthEnv | Tuanmu & Jetz 2014 |
| Heterogeneity (0.0-1.0) | EarthEnv | Tuanmu & Jetz 2015 |
| Mosaic forest (%) | EarthEnv | Tuanmu & Jetz 2014 |
| Terrain Roughness Index (TRI) | ENVIREM | Title & Bemmels 2018 |

Heterogeneity is a biophysical similarity measure used here to define canopy species richness (i.e., canopy structure, composition and diversity) derived from textural features of Enhanced Vegetation Index (EVI) between adjacent pixels; sourced from the Moderate Resolution Imaging Spectroradiometer (MODIS, https://modis.gsfc.nasa.gov/). We inverted the raster cell values in the original EarthEnv variable ‘Homogeneity’ (Tuanmu & Jetz 2015) to represent the spatial variability and arrangement of canopy species richness on a continuous scale which varies between zero (minimum heterogeneity, low species richness) and one (maximum heterogeneity, high species richness). The three measures of percentage landcover (Evergreen Forest, Mosaic Forest, Cultivated) are consensus products integrating GlobCover (v2.2), MODIS land-cover product (v051), GLC2000 (v1.1) and DISCover (v2) at 30 arc-sec (~1-km) spatial resolution. Mosaic forest is derived from the EarthEnv variable ‘Mixed trees’ and represents a mosaic of mixed forest, shrubland and woody savanna, with cultivated representing a mix of cropland, tree cover and managed vegetation. Full details on methodology and image processing can be found in Tuanmu & Jetz (2014) for the landcover layers, and Tuanmu & Jetz (2015) for the habitat heterogeneity texture measure.

After selecting biologically relevant covariates, all six candidate variables were tested for multicollinearity using Variance Inflation Factor (VIF) analysis in the R package usdm (Naimi *et al*. 2014). VIF is based on the square of multiple correlation coefficients, regressing a single covariate against all other covariates. A stepwise elimination of highly correlated variables was used retaining covariates with a VIF threshold of < 5, considered as suitable for multi-variable correlation (Dormann *et al*. 2013). The remaining covariates were then checked for collinearity using Spearman’s Correlation Coefficient with a threshold of *r_s_* = |0.7|. All the selected covariates showed low collinearity and thus all were included as predictors in model calibration (Table S1).

### Resource Selection Functions

We calibrated RSFs for each species using logistic regression fitted as generalised linear models (GLMs) with a binomial logit link function in the ENMTools (Warren *et al*. 2019b) and stats R packages (R Core Team, 2018). Our RSFs followed geographical range first-order selection (Johnson 1980) using design I in a use-availability sampling protocol (Manly *et al*. 2002; Johnson *et al*. 2006; Thomas & Taylor 2006). GLMs were fitted with iteratively reweighted least squares to derive maximum likelihood estimates on model parameters, with no interaction terms. Linear and quadratic terms were fitted dependent on the scaled responses from fitting both terms on an initial model. Only linear terms were used when the quadratic term resulted in biologically unrealistic U-shaped curves, or when a linear term was sufficient to explain the scaled response. All covariates were standardized with a mean of zero and standard deviation of one.

Background-absence points were randomly sampled using 10,000 points suitable for regression-based modelling (Barbet-Massin *et al*. 2012) and weights assigned equally to both presence and background points. We tested calibration accuracy using the explained variance from each logistic model measured using McFadden’s adjusted *R^2^* (*R^2^_adj_*, McFadden 1974). We tested discrimination ability in environmental space, using a new implementation of the AUC (Area Under the Curve) statistic (referred to here as AUC_env_) calculated using the ‘env.evaluate’ function in the R package ENMTools (Warren *et al*. 2019b). AUC_env_ uses Latin hypercube sampling to estimate model fit in environmental space using all possible environmental combinations, evaluating model performance between presence and background-absence points within the minima and maxima of the environmental predictors. An AUC_env_ = 1.0 indicates maximum predictive model performance, with an AUC_env_ = 0.5 being no better than a random prediction.

Given the inherent spatial autocorrelation and environmental heterogeneity present in biodiversity inventory data, we used a Monte-Carlo randomization to test for significance in the AUC evaluation metrics against null random expectations (Raes & ter Steege 2007). We set random null models to calculate 95 % AUC Confidence Intervals (CI) on a frequency histogram, randomly drawing points without replacement on 100 replicates using 20 % test data under the same environmental parameters used in the best-fit models. With fitted 95 % CI AUC values, model accuracy was assessed on being higher than expected by chance at *α* = 0.05. Marginal response functions were used to assess habitat differences between both eagle species and biological realism of the fitted models (Guevara *et al*. 2017). Marginal responses show the response of an RSF to the habitat covariate with all other covariates held at their mean in the available environment.

### Resource overlap

#### RSFs

We quantified the level of resource specialization for both eagles, using Levins (1968) standardized niche breadth *B2* statistic for each GLM in geographic (*gB2*) and environmental (*eB2*) space given a vector of habitat suitability scores. Niche breadths were measured on a scale from 0 to 1, with zero indicating low niche breadth (habitat specialist) and 1 high niche breadth (habitat generalist; Krebs 1999). Second, we calculated pair-wise niche overlap metrics on each respective GLM in both geographic and environmental space to quantify similarity between models using Schoener’s *D* (Schoener 1968, Warren *et al*. 2008). Schoener’s *D* measures niche similarity between environmental conditions ranging from 0 (no overlap) to 1 (identical predictions). Niche breadth and overlap measures were calculated based on Monte Carlo integration using continuous predictions in the R package ENMTools (Warren *et al*. 2019a,b).

#### PCA-env ordination

For the PCA-env, we included the same six covariates (Table 1) at a spatial resolution of 2.5 arc-minutes (~4.5-km resolution). All 30 arc-sec (~1-km resolution) landcover layers were thus resampled to a spatial resolution of 2.5 arc-minutes using bilinear interpolation. We thinned occurrences using a 5 km spatial filter to approximately match the resolution of the raster data (~4.5-km). Applying the 5-km spatial filter resulted in 353 occurrences for the crested eagle and 606 occurrences for the harpy eagle. The 5-km filtered occurrences for both eagle species had a random and independent distribution (*O* = 0.525) compared to the unfiltered occurrences.

Ordination techniques such as principal component analysis (PCA) use direct comparisons of species-habitat relationships in environmental space in contrast to predictive resource overlap analysis using RSFs. Using the PCA-env framework of Broennimann *et al*. (2011), we performed an environmental ordination using the R package humboldt (Brown & Carnaval 2019). We set the PCA-env method to calibrate on a non-analogous environmental space using a minimum convex polygon around all spatially filtered occurrences on a 100 x 100 resolution grid, with a smoothed Gaussian kernel density function (bandwidth = 1) to account for spatial auto-correlation. We quantified niche overlap on the first two principal components using Schoener’s *D* statistic. Using smoothed densities allows measured overlap to be independent of grid resolution, important for unbiased estimates of niche overlap using Schoener’s *D* (Broennimann *et al*. 2011).

To test equivalency in shared environmental space, first, we used a niche Equivalency Statistic to test for the difference (*α* = 0.05) between the observed overlap scores and those under a null distribution hypothesis that the two distributions are equivalent (Warren *et al*. 2008). For the null distribution, presence points are randomly assigned to each species, and a PCA is built on these randomized data. This is repeated a hundred times and a probability distribution is then estimated for niche overlap under the null hypothesis that both sets of species occurrences are randomly distributed in the environment. Second, to measure the ability of the Equivalence Statistic to detect differences in environmental space, we used a Background Statistic to test for the difference (*α* = 0.05) if the observed occurrences of one species are more similar than expected by chance to the background occurrences of the other species (*n* = 100; Warren *et al*. 2008).

The background test corrects for the environmental heterogeneity inherent in environmental data underlying occurrence data, assuming that all species are choosing environments at random throughout their respective geographic ranges (Warren *et al*. 2008). If distributions are not equivalent, a statistically significant difference allows rejection of the null hypothesis of niche equivalency between the two distributions, regardless of the significance of the Background Statistic. A non-significant Equivalence Statistic and significant Background Statistic supports the hypothesis of equivalent shared environmental space. If both statistics are non-significant this implies niche equivalency could be the result of shared identical environmental space, with limited power for the Equivalency Statistic to detect any significant differences (Brown & Carnaval 2019). Importantly, the Background Statistic assesses the power of the equivalency test by asking if two distribution models are equivalent based on the matching environments available. It does not provide any evidence that niches are not equivalent. General model development and geospatial analysis were performed in R (v3.5.1; R Core Team, 2018) using the dismo (Hijmans *et al*. 2017), raster (Hijmans 2017), rgdal (Bivand *et al*. 2019), rgeos (Bivand & Rundle 2019) and sp (Bivand *et al*. 2013) packages.

## Results

### Resource Selection Functions

Both GLMs had high discrimination ability (crested eagle: AUC_env_ = 0.984, harpy eagle = AUC_env_ = 0.976), with randomization tests robust against null expectations (*p* < 0.05; Fig. S1). For the crested eagle, seven covariates had significant terms (Table S2), with the full model able to explain 88 % of the variability in environmental space (*R^2^_adj_* = 0.88). Crested eagles were more likely to be positively associated with high canopy species richness and evergreen forest, but negatively associated with mosaic forest cover comprising mixed forest and woody savanna, elevation, and cultivated areas. For the harpy eagle, seven covariates had significant terms (Table S2), with the full model able to best explain 86 % of the variability in environmental space (*R^2^_adj_* = 0.86). Harpy eagles were more likely to be positively associated with evergreen forest, and negatively associated with elevation, with responses to both these covariates stronger compared to the crested eagle. The harpy eagle had similar negative associations with cultivated areas and mosaic forest cover, as the crested eagle.

Overall, responses in distribution to habitat were consistent for both the crested and harpy eagle, with similar responses to canopy species richness and topography and restricted tolerances for higher terrain roughness (Fig. 2). Both eagles were positively associated with areas of high heterogeneity, indicating similar preference for highly species-rich canopy vegetation. Responses to the three landcover covariates, however, differ subtly. Crested eagles had high peak suitability of 60 % evergreen forest cover, compared with the higher response higher for the harpy eagle at 75 % evergreen forest cover, suggesting greater reliance on this landcover type for the harpy eagle. For mosaic forest cover, the crested eagle showed a steeper negative response to higher percent mixed tree and savanna landcover, compared to the weaker negative response from the harpy eagle. For cultivated areas, both eagles had fairly broad tolerances but with highest suitability in areas with 20 % cultivation cover.

**Figure 2.**
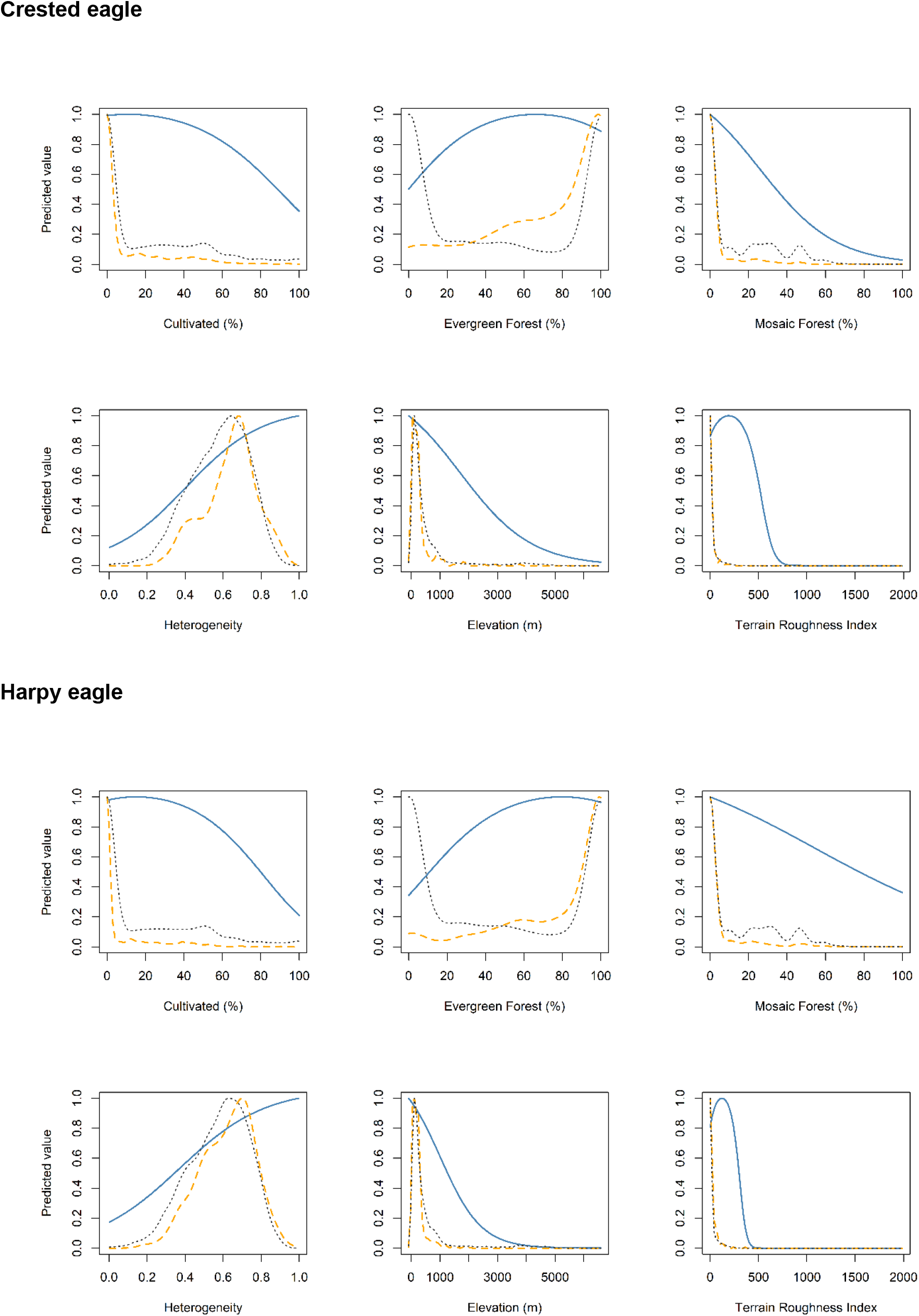
Marginal responses for each respective species model from all continuous GLM covariates. Blue lines indicate environmental suitability, orange dashed line presence locations and grey dotted line background locations.

### Environmental overlap

From the GLMs both eagles had similar niche breadth across geographical space, with crested eagle niche breadth marginally higher (*gB*2 = 0.679), than the harpy eagle (*gB*2 = 0.650). In environmental space, niche breadth was markedly reduced for both the crested eagle (*eB*2 = 0.034) and harpy eagle (*eB*2 = 0.035), meaning that both eagles have specialized habitat requirements. Resource overlap across geographical space from the GLMs was high (*D* = 0.926), but moderate across environmental space (*D* = 0.665). Measuring overlap in environmental space using the coarser-grain PCA-env resulted in moderate overlap (*D* = 0.762, Fig. 4), with both eagles sharing equivalent environmental space supported from the combination of a non-significant Equivalency Statistic (*p* = 0.79) and significant Background Statistics (*p* = 0.01). Both eagles occupied more similar environmental space than expected by chance, with the harpy eagle occupying a slightly more restricted environmental space compared to the crested eagle (Fig. 5).

**Figure 4.**
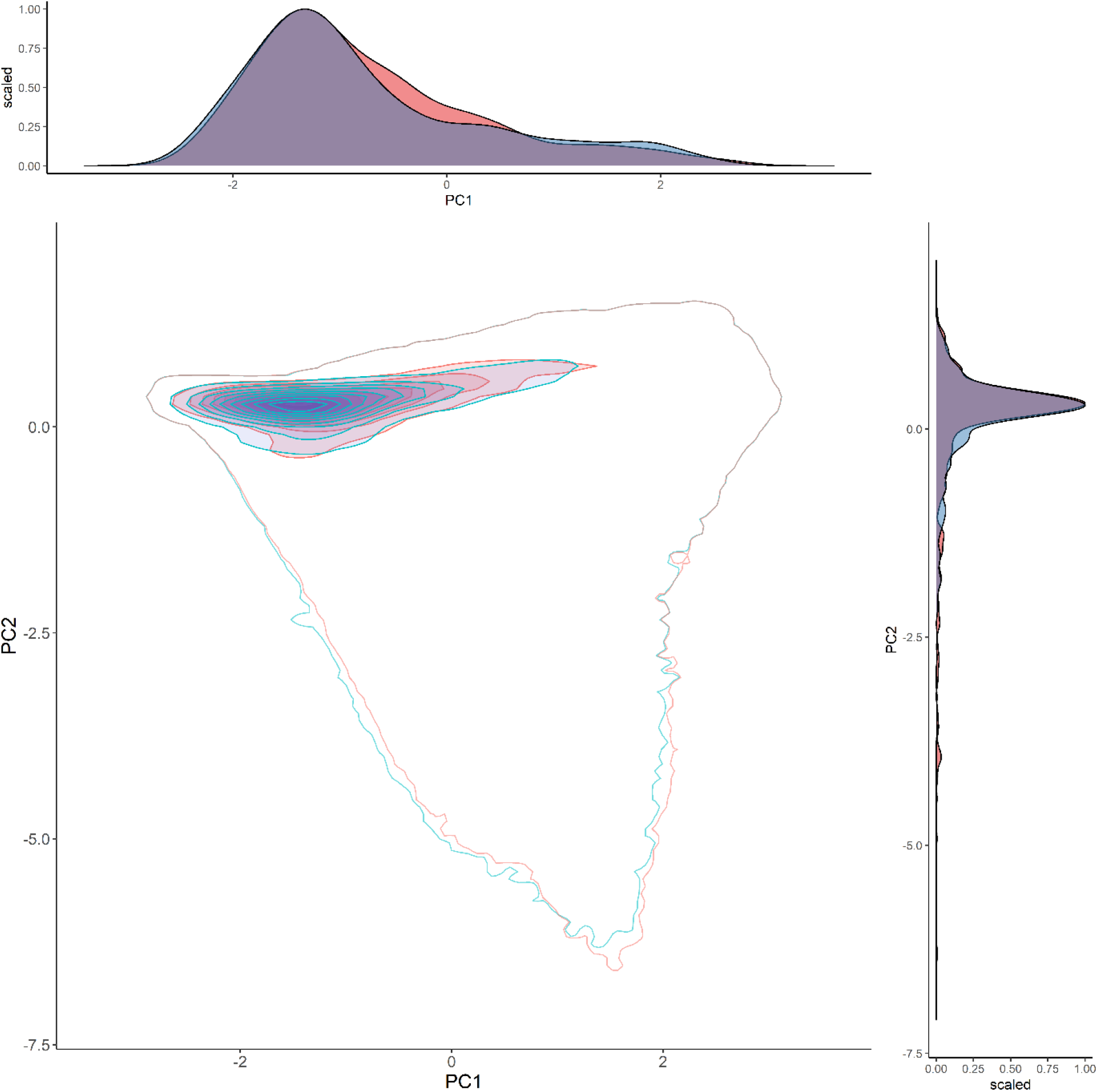
Environmental overlap (purple) for the crested eagle (red) and harpy eagle (blue) across environmental space from two principal components. Total variance explained by the two principal components = 73.34 % (PC1 = 45.81 %, PC2 = 27.53 %). Filled isopleths are kernel densities from 1-100%. Empty kernel density isopleths represent 1% density isopleth of the environment.

**Figure 5.**
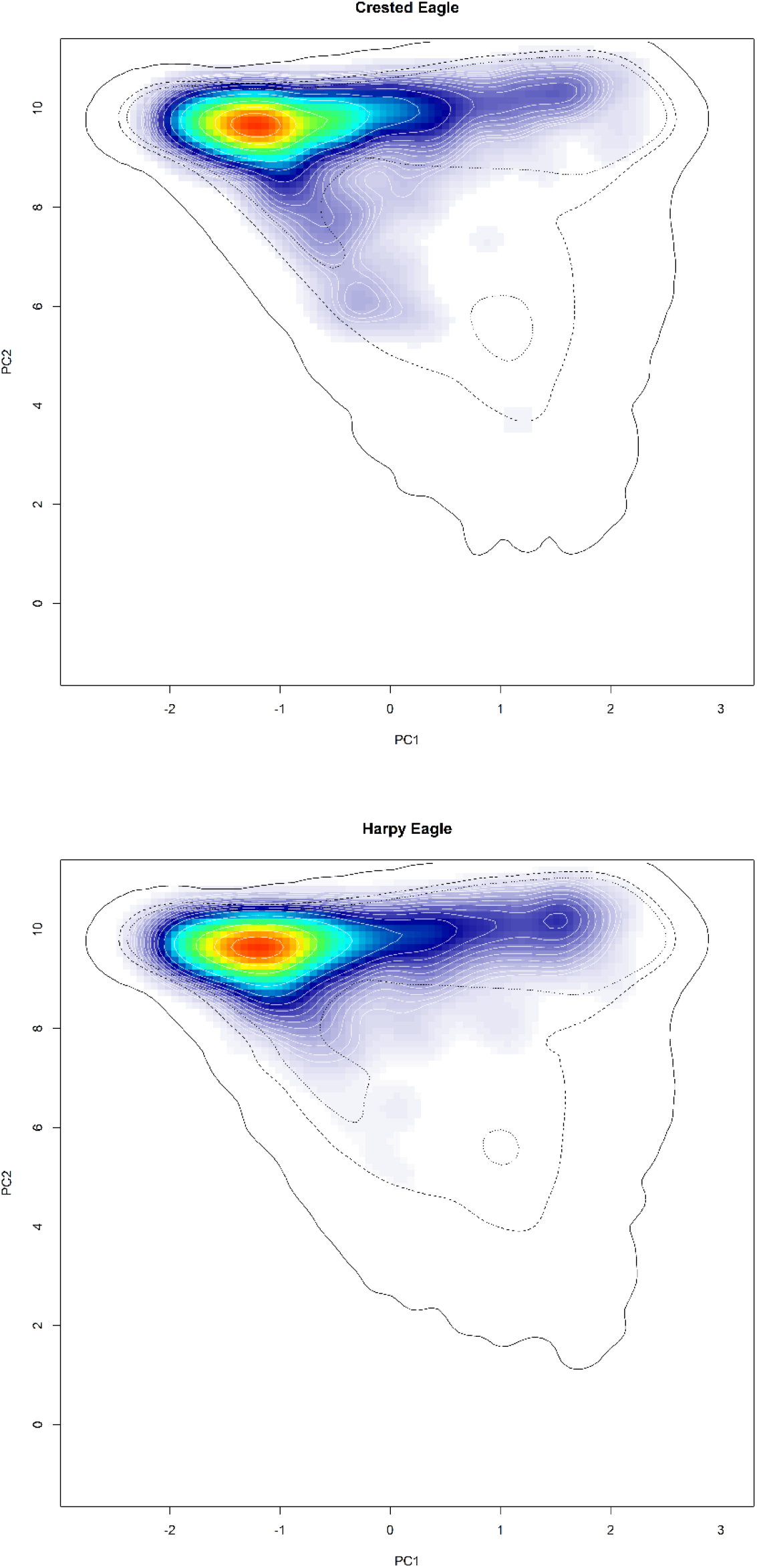
Distribution in environmental space for both eagles across two principal components. Red areas indicate highest environmental suitability. Filled kernel density isopleths characterize kernel density values from 0.4 (blue) to 0.99 (red). Black isopleth lines define kernel density of the corresponding environment, with black solid line = 0.1, black hashed line = 0.5, black dotted line = 0.75.

## Discussion

Previous localized studies on the co-occurrence of the crested and harpy eagle have produced contrasting results on the level of shared environmental space (Thiollay 1989; Sanaiotti *et al*. 2015). Here, we used a biogeographical perspective to identify environmental differences between these two eagles, knowing that factors affecting distribution in certain areas may not be generalizable into other regions within their shared range. Our predictive GLMs identified narrow environmental niche breadths for both eagles, despite broad niche breadth measured in geographical space. Whilst resource overlap in geographical space was high using GLMs, there was only moderate niche overlap in environmental space using both GLMs and PCA-env. Both eagles shared identical environmental niches from randomization tests, with only subtle differences in distribution across environmental space. Thus, at the biogeographic scale both species can co-exist, presumably if there is sufficient habitat heterogeneity for both eagles to coincide when they occur in close proximity (see Tilman 2004).

### Environmental differences

Most species are expected to be non-randomly distributed in environmental space and to occur within an optimal range of conditions (Hirzel *et al*. 2002). For the crested and harpy eagle, seven primary limiting factors were sufficient to explain both their distributions from the GLMs. Both distributions were positively associated with high canopy heterogeneity, indicating a preference for areas of highly species-rich canopy vegetation and structural complexity, although the association was stronger for crested eagle than harpy eagle. Diverse species richness and structural diversity of canopy trees may offer greater diversity of prey species (Tews *et al*. 2014; Stein *et al*. 2014) and also attack methods to the dietary generalist crested eagle. Harpy eagle, in contrast, are more specialized in diet and likely hunting strategy, and may benefit from a simpler forest structure where prey detection and ambush are more ritualized for the larger and presumably less agile specialist harpy eagle.

However, the harpy eagle was more negatively associated with elevation and TRI compared to the crested eagle. This suggests that the harpy eagle is more restricted by topography than the crested eagle, especially in topographically complex areas, such as Darien in Panama, compared to vast, low-lying areas across the Amazon basin. Perhaps nesting in emergent trees requires harpy eagles to select flatter, lower elevation areas where these tall trees are more common (Miranda *et al*. 2020; Vargas González *et al*. 2020) and with the required T-shaped architecture to build the large nest (Pinho *et al*. 2020). Further, both eagles had positive responses to increased evergreen forest cover (Fig. 2), but with the crested eagle less likely to be associated with this vegetation type than the harpy eagle (Table S2). Thus, the crested eagle may inhabit areas with less evergreen forest cover, partly explaining how the two species co-exist – with crested eagles possibly able to persist into drier seasonal forest environments (Whitacre *et al*. 2012).

Although both eagles were most likely to be associated with areas of abundant evergreen forest, there were only slight differences in response to suitable landcover types. Both eagles had negative responses to mosaic forest cover, but with the crested eagle more strongly negatively associated with them than the harpy eagle. Further, both the crested and harpy eagle had a broad tolerance to cultivated landcover, though highest suitability occurred in areas of < 20 % cultivated landcover. This suggests that both eagles may struggle to adapt to more fragmented cultivated habitat, particularly for the harpy eagle which does not switch to terrestrial prey in cultivated areas, resulting in a lack of food provisioning for nestlings (Miranda 2021). Conversely, the crested eagle may be more reliant on mature forest, possibly associated with their stronger reliance on a diet of arboreal snakes and cavity nesting arboreal birds and mammals (Whitacre *et al*. 2012; Gomes *et al*. 2021). Even so, this does not consider how this may affect breeding performance, with harpy eagles having lower breeding productivity in fragmented habitats (Miranda *et al*. 2021).

### Resource overlap

Measuring the level of resource overlap in both geographical and environmental dimensions further revealed how both eagles respond to habitat conditions at different spatial resolutions. Measuring overlap in geographical space is relevant only for models solely estimating distribution. Whereas, measuring resource overlap in environmental space will often reveal the underlying processes determining the occupation of differing environmental space (Warren *et al*. 2019a). Our results support differences between geographical and environmental space, demonstrating the importance of measuring resource overlap across both geographic and environmental dimensions. Further, altering the spatial resolution had little effect on the general pattern of moderate resource overlap. Importantly, both eagles had narrow niche breadth in environmental space when compared to geographic space, indicating habitat specialization. This follows the general observation for both eagles being generally reliant on highly speciose primary tropical forest (Whitacre *et al*. 2012; Miranda *et al*. 2019).

Using ordination to directly measure species-habitat relationships at a coarser grain showed a similar moderate resource overlap to the fine-grain GLMs. Randomization tests showed no support for differing spatial niches, demonstrating that both eagles generally inhabit similar environmental spaces within the same geographical range. This concurs with the theory of niche conservatism, where the expectation is for greater niche equivalence between pairs of sister-species (Wiens & Graham 2005; Warren *et al*. 2008). Niches can be conserved over long evolutionary timescales, with sister-species likely to inhabit similar, though not identical environments (Wiens & Graham 2005). Given the high genetic similarity (~91 %) between these two eagle species (Lerner & Mindell 2005), the moderate resource overlap and equivalent niches supporting niche conservatism were expected. Thus, our results show no support for competition restricting geographic range limits between either species, when regarding vegetation heterogeneity and topography. However, differing landcover use may be driven by the specific diet preferences of both eagles. Thus, any differences in habitat resource use may be occurring at a finer micro-habitat scale, which are undetectable at the continental geographic scale used here. Future research should thus focus on habitat use at finer scales, for example testing for differences in nesting elevation. Harpy eagles generally nest below 350 m/asl (Vargas González & Vargas 2011; Vargas González *et al*. 2020) but crested eagles are known to breed at higher elevations (Gomes & Sanaiotti 2015).

Determining overlap in habitat use between pairs of potentially competing species has long been a focus in ecology (Levins 1968; MacArthur 1972). Using logistic regression and ordination has been effective here for identifying overlap in both distribution and aspects of the habitat that can determine the nature and abundance of available resources between these two tropical forest raptors. Both the crested and harpy eagle occupy equivalent environmental space within similar geographical ranges and show a moderate (but not significantly different) level of habitat overlap. Because both eagles share the same environmental space, coexistence is likely driven by the availability of each eagle’s specific food resources, for which the overlap is minimal. Habitat preference may be driven by where the most favourable food resources are distributed, perhaps accounting for the subtle differences in landcover preference within their shared range. Our results highlight the importance of measuring geographical and environmental overlap using multiple grain resolutions to gain a broad understanding of the processes determining distribution for co-existing species with similar geographical ranges. Smaller, individual-scale studies of resource partitioning are necessary to further clarify the determinants of their co-existence.

## Data Accessibility Statement

Upon acceptance the data that support the findings of this study will be made openly available on the data repository *figshare*

## Conflict of Interest statement

The authors have no conflicts of interest to declare.

## Appendices

### Appendix 1 Supplementary Tables

**Table S1.**
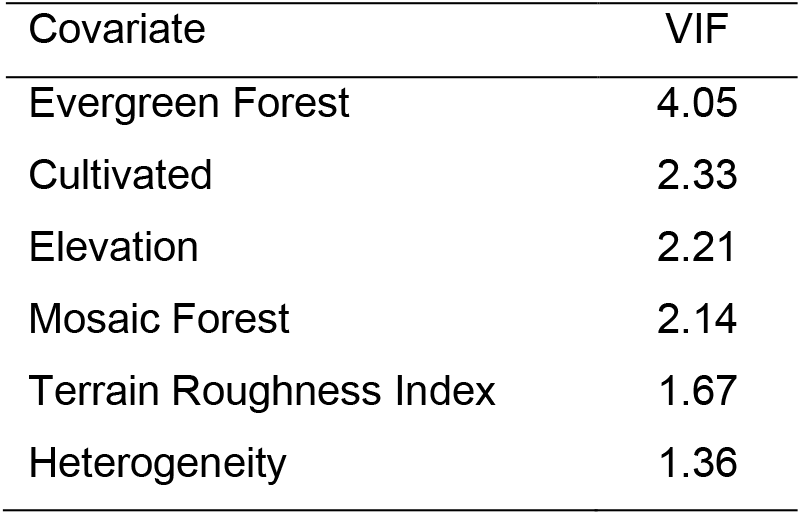
Multi-collinearity test using stepwise elimination Variance Inflation Factor (VIF) analysis for the GLMs. Covariates with VIF < 10 have low correlation with other variables, and thus are suitable for inclusion in calibration models when further evaluated for ecological relevance.

| Covariate | VIF |
| --- | --- |
| Evergreen Forest | 4.05 |
| Cultivated | 2.33 |
| Elevation | 2.21 |
| Mosaic Forest | 2.14 |
| Terrain Roughness Index | 1.67 |
| Heterogeneity | 1.36 |

**Table S2.** GLM terms derived from maximum likelihood estimates obtained from each respective species model. Covariates ranked by the value of regression coefficient estimates. Superscript 2 indicates quadratic model terms.

| Parameter | Estimate | SE | *z* | *p* |
| --- | --- | --- | --- | --- |
| Intercept | -0.83 | 0.12 | -6.70 | <0.001 |
| Heterogeneity | 79.89 | 11.85 | 6.74 | <0.001 |
| Mosaic Forest | -79.40 | 18.12 | -4.38 | <0.001 |
| Elevation | -63.36 | 22.99 | -2.76 | 0.001 |
| Evergreen Forest | 55.08 | 21.36 | 2.58 | 0.001 |
| Cultivated | -36.43 | 17.86 | -2.04 | 0.041 |
| Cultivated<sup>2</sup> | -29.93 | 15.28 | -1.96 | 0.050 |
| Evergreen Forest<sup>2</sup> | -28.60 | 9.84 | -2.91 | 0.004 |
| Terrain Roughness Index<sup>2</sup> | -14.35 | 14.42 | -0.10 | 0.320 |
| Terrain Roughness Index | 5.68 | 15.35 | 0.37 | 0.711 |

Harpy eagle
| Parameter | Estimate | SE | *z* | *p* |
| --- | --- | --- | --- | --- |
| Intercept | -0.96 | 0.11 | -8.87 | <0.001 |
| Elevation | -135.71 | 26.35 | -5.15 | <0.001 |
| Evergreen Forest | 82.25 | 18.24 | 4.51 | <0.001 |
| Heterogeneity | 68.81 | 8.63 | 7.97 | <0.001 |
| Mosaic Forest | -41.85 | 14.38 | -2.91 | 0.004 |
| Cultivated | -41.51 | 15.39 | -2.70 | 0.007 |
| Evergreen Forest<sup>2</sup> | -31.76 | 7.74 | -4.10 | <0.001 |
| Terrain Roughness Index | 27.10 | 12.62 | 2.15 | 0.031 |
| Cultivated<sup>2</sup> | -19.64 | 13.08 | -1.50 | 0.133 |
| Terrain Roughness Index<sup>2</sup> | -13.21 | 8.27 | -1.60 | 0.110 |

### Appendix 2 Supplementary Figures

**Figure S1.**
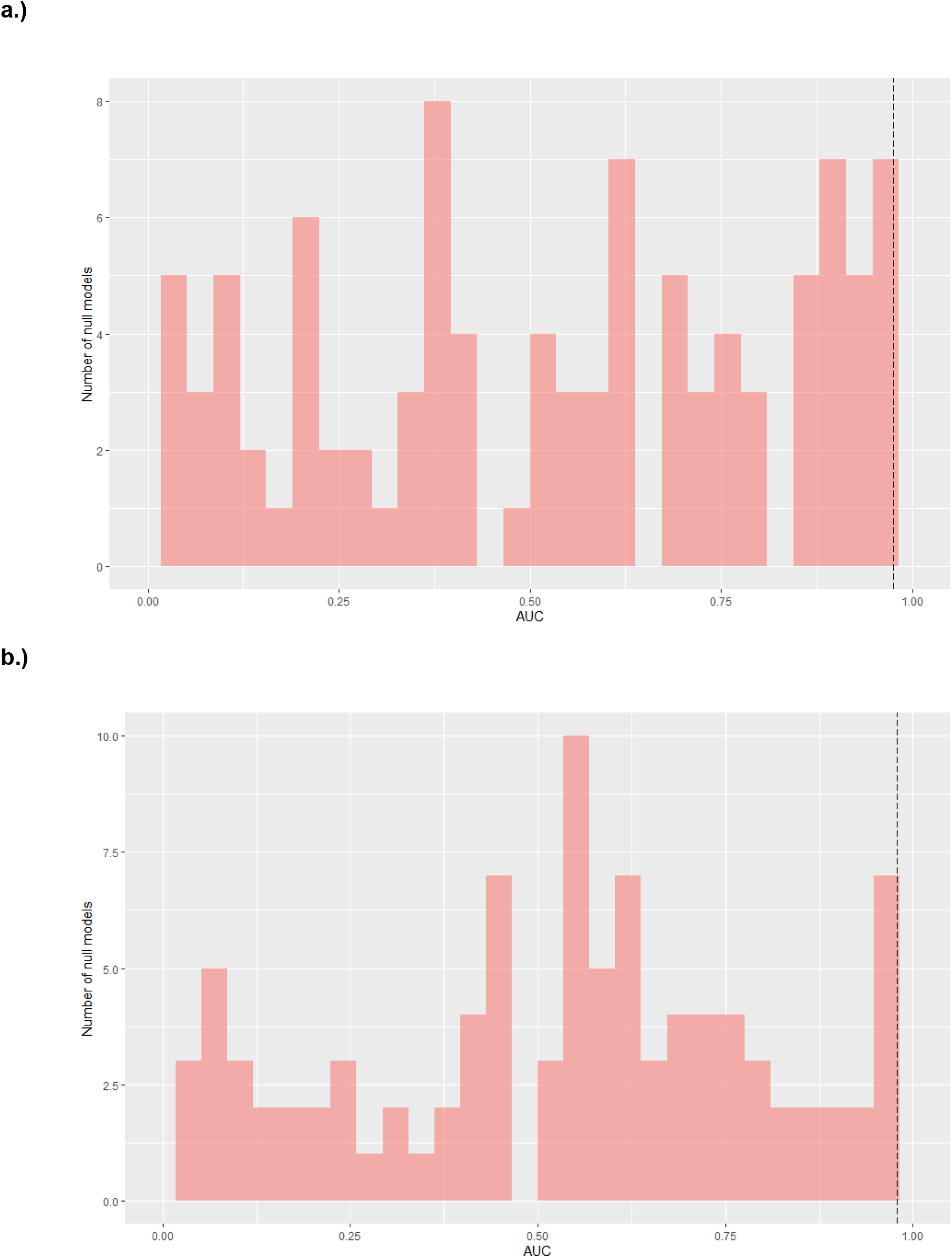
Monte-Carlo randomization test based on 100 random null models to test significance against best-fit model for the crested eagle (a) and harpy eagle (b) in environmental space. Dashed vertical line indicates AUC for each best fit model.

## Notes

### Competing Interest Statement

The authors have declared no competing interest.

